# Are the Matrix Metalloproteinases 2 and 9 beeing expressed in the mucosa of the rat small intestine during its intrauterine and postnatal life development ?

**DOI:** 10.1101/619767

**Authors:** Camila Audrey dos Reis, José Rosa Gomes

## Abstract

MMP-2 and MMP-9 are proteins with well stablished roles on the remodeling of tissue during morphogenesis. This study aimed to evaluate the activity and expression of the MMP-2 and MMP-9 in the rat small intestine mucosa layer on 15^th^ and 18^th^ days of intratuterine life (i.u.) and at 3rd, 10^th^, 17^th^, 25^th^, and 32^th^ days after birth (a.b.). Samples were submitted to zimography, immunohistochemistry methods and Masson trichrome staining. Results showed that MMP-2 and MMP-9 were not expressed in the small intestine epithelium during intrauterine life. MMP-9 was immunolocalized in the villi goblet cells and in the lamina propria in cells identified as being the mast cells (a.b.). We concluded that in the i.u. and a.b. life the MMP-2 and MMP-9 were not expressed in the small intestine epithelium. However, after birth, because MMP-9 was expressed in the mast cels present in the lamina propria it may be involved in the remodeling process of the innate immunity that occurs during the small intestine development.

## INTRODUCTION

In the mammalian, small and large intestines begin their development from the lateral fusion of the endoderm (Noah et al, 2011) during the intrauterine (i.u.) life. In rats, the primordial digestory tube formed is lined by undifferentiated epithelial cells arranged in multilayers that are maintained until approximately the 13th day of i.u. life (Smith et al., 2000; Louvard et al., 1992), where the small intestine differentiation processes begins. Around the 19th day (i.u.) it is already possible to detect villi structures emerging from the epithelium surface. After birth, rat pass through suckling and weaning phases, in which several changes occur in the intestinal epithelium to ensure the differentiation of the various cell types such as endocrine and goblet cells as well as the closure of the absortive process.

During the suckling and weaning phases, the intestinal epithelium becomes completely developed, and two compartments can be identified; the villi (functional compartment) and the crypts, which are considered the proliferative compartment where cells are produced and differentiated, before their migration to the villi to execute their digestive functions.

The suckling phase represents the period from birth until the 16th day of life, where milk is the main food. Around the 17th day, during the third week of life, diet with milk is changed to diet with solid food (weaning phase). Therefore, during the first three weeks after birth, the mucosa layer, specially the epithelium, goes through several morphological and physiological remodeling related to digestive and absorptive processes. These functional changes are produced by a shift in the expression of different enzymes present in the cell membrane like lactase, maltase, sucrase and intestinal alkaline phosphatase (Goldstein et al, 1971; Uezato and Fujita, 1983;Tojyo, 1984; Gomes et al., 2016).

However, in this decade it has been demonstrated that another kind of enzymes denominated matrix metalloproteinases seem to have important roles during the development, growth and morphological maturation of different parts of the body like bones (Holmbeck et al, 1999; Yang et al., 2008), vascular system, skeletal muscle (Oh et al., 2004s), kidney and dental structures (Gomes et al., 2010, 2011, Omar et al., 2011, 2017).

Metalloproteinases found in the extracellular matrix are denominated matrix metalloproteinases (MMPs) such as MMP-2 and MMP-9 (also denominated gelatinases) or may be anchored in the cell membrane which is denominated membrane matrix metalloproteinase (MT-MMPs). Despite this, there are still few descriptions about the involvement of MMPs in this process concerning to the development of the small intestine (Kurakata et al., 2008; Camargo et al., 2016) and none report regarding expression, activity or functions for the gelatinases MMP-2 and MMP-9.

Since rat small intestine has a continuous morphological and physiological remodeling process along its development, we hypothesized that MMP-2 and MMP-9 could present a relevant role during its embryonic and postnatal development. As such, this investigation aimed to evaluate MMP-2 and MMP-9 expression in the small intestine of rats, from the intrauterine (15th day) to post-natal life (32th day) of life. Our results pointed to a possible role only for MMP-9 regarding to the development of the small intestine development.

## MATERIAL AND METHODS

The university’s Committee for Ethics in Animal Research approved all experiments in the study with the protocol number 12790/2010.

### Animals

Adult female and male Wistar rats from the State University of Ponta Grossa colony were mated and the vaginal fluid was collected and spread on a slide to detect the presence of spermatozoids for embryo sampling at 15^th^ and 18^th^ days of intrauterine life. Other pregnant rats were separated to obtain the pups evaluated after birth on the 3rd, 10^th^, 17^th^, 25^th^ days and 32^th^ days. All rats were maintained under conventional conditions with a 12h light/dark cycle (lights on at 06:30h/lights off at 18:30h) at 25°C and received food (balanced ration from Nuvital, Brazil) and water *ad libitum*.

### Embryonic and Postnatal Times Evaluated

Females were briefly anesthetized with **Ketamine** 100 mg/kg (Laboratório Cristália, Itapira-SP) and **Xylazine** 10mg/kg (**ROMPUM** - **Bayer** do **Brasil** S/A - São Paulo - S.P.) and killed by cervical dislocation. Six embryos were sampled on the 15^th^ and 18^th^ days of intrauterine life, and six male rats were used for small intestine sampling the after birth (a.b.), on the 3rd, 10^th^, 17^th^, 25^th^ days and 32^th^ days. Three embryos and rats (a.b.) were used for zymography method and three for the histology procedures.

### Zymography and MMP inhibitor test

Fragments of small intestine collected were homogenized in 500 µl of ice buffer (50 mM Tris-HCL pH 7.5 containing 5mM CaCl and 0.9% NaCl). The resulting homogenate was then briefly sonicated (3x 50s), cleared by centrifugation at 15,000g for 15 min at 4°C and pooled with equal volume from each sample. Each of sample of small intestine was submitted to total protein quantification by the Bradford method (1976) and to zymography, which demonstrated the same pattern of the results for each sample (data not showed). Then we decided to produce a pool of the samples by time evaluated. For zymography, the following procedures were perfomed: from each pool, 0,1 µg of total protein were diluted in non-reduced Laemmli buffer and were submitted to electrophoresis in a 10% polyacrylamide gel containing 5% gelatin, for 4 h. Gels were rinsed twice (30min) in 2.5% Triton X-100, under shaking, and then immersed in activation buffer (2M CaCl_2_; 1M HCl) for 16 h at 37°C. Gels were stained with Coomassie Blue until the degraded bands could be seen. Other gels were submitted to inhibition tests using 1,10-fenantrolina e EDTA 5mM (MMP inhibitors); iodoacetamide 1mM (cisteine inhibidor) and phenylmethylsulfonyl fluoride 2mM (PMSF-serine inhibitor). The inhibitors were purchade from SIGMA: PMSF cod. 329-98-6; EDTA cod. 60-00-4; PHE cod 66-71-7; Iodoacetamide cod 144-48-9

### Histological procedures

For each of the intrauterine times, a cross section in each embryo was performed at the thoracic level and the abdominal part. Sections were immersed in 2% formaldehyde prepared in 0.1M phosphate buffer, pH 7.4, for 48 hours. After birth, the jejunum fragments were submitted to the same fixative solution for the same time. Afterwards, samples were dehydrated in alcohol, rinsed in xylol and embedded in paraffin to obtain three semi-seriated sections of 5µm. The sections were mounted on slides for immunohistochemical procedures.

### Immunohistochemistry for MMP-9 and MMP-2

For this analysis, three slides containg three embryon, and three fragment of small intestine (obtained after birth) were dewaxing and submitted to antingenic recovery in EDTA 1mM pH 8.0 at 98°C for 20 min. After that, sections were quenched three times with 2% hydrogen peroxide for 10 min each to inhibit endogenous peroxidase activity. Sections were washed in water and PBS pH 7.4 and immunostained with primary antibodies (2ug/ml, purchased from Millipore USA) as follows: MMP-9 (1:250 cod MAB3309 and MMP-2 (1:250- cod MAB3308) in PBS containing 1% BSA, and incubated overnight. Sections were then washed in PBS and incubated with secondary antibody using DAKO (Universal LSAB kit Cod K067511-2) for 30 min at 37C in each solution. After washing in PBS, sections were incubated with the DAB reagent (SIGMA - cod D3939) prepared in PBS in the presence of hydrogen peroxide. Negative control staining was performed for each molecule by either omitting the primary antiserum or substituting the primary antiserum with nonimmune serum.

## RESULTS

### MMP-9 and MMP-2 were not expressed during the development of small intestine in the intrauterine life

According to zymograms results, demonstrated in the figure 1 A, there is no proteolytic activity for gelatin degradation at 15th and 18th days of the intrauterine life. These results were confirmed by the immunohistochemistry presented in Figure 2 A, B, C, and D for the times evaluated, in which the MMP-2 and MMP-9 were not detected.

**Fig. 1.**
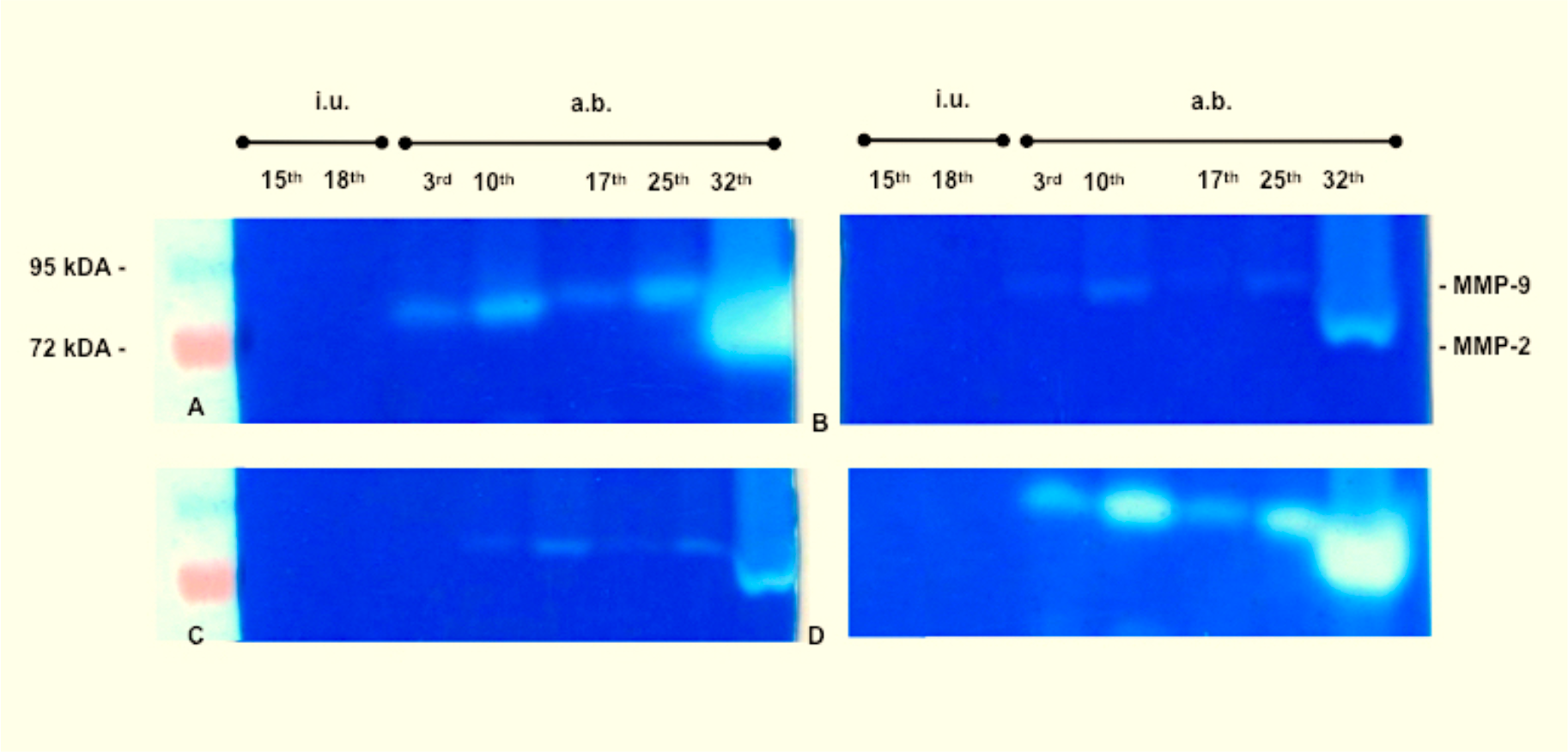
Representative Zymography (A) and inhibitory tests (B,C and D) using 0.5 mM of PHE + EDTA; 2mM of PMSF and 1mM of iodocetamide for MMPs, Serinases and Cysteinases respectively along of the times of small intestine development.

**Fig. 2.**
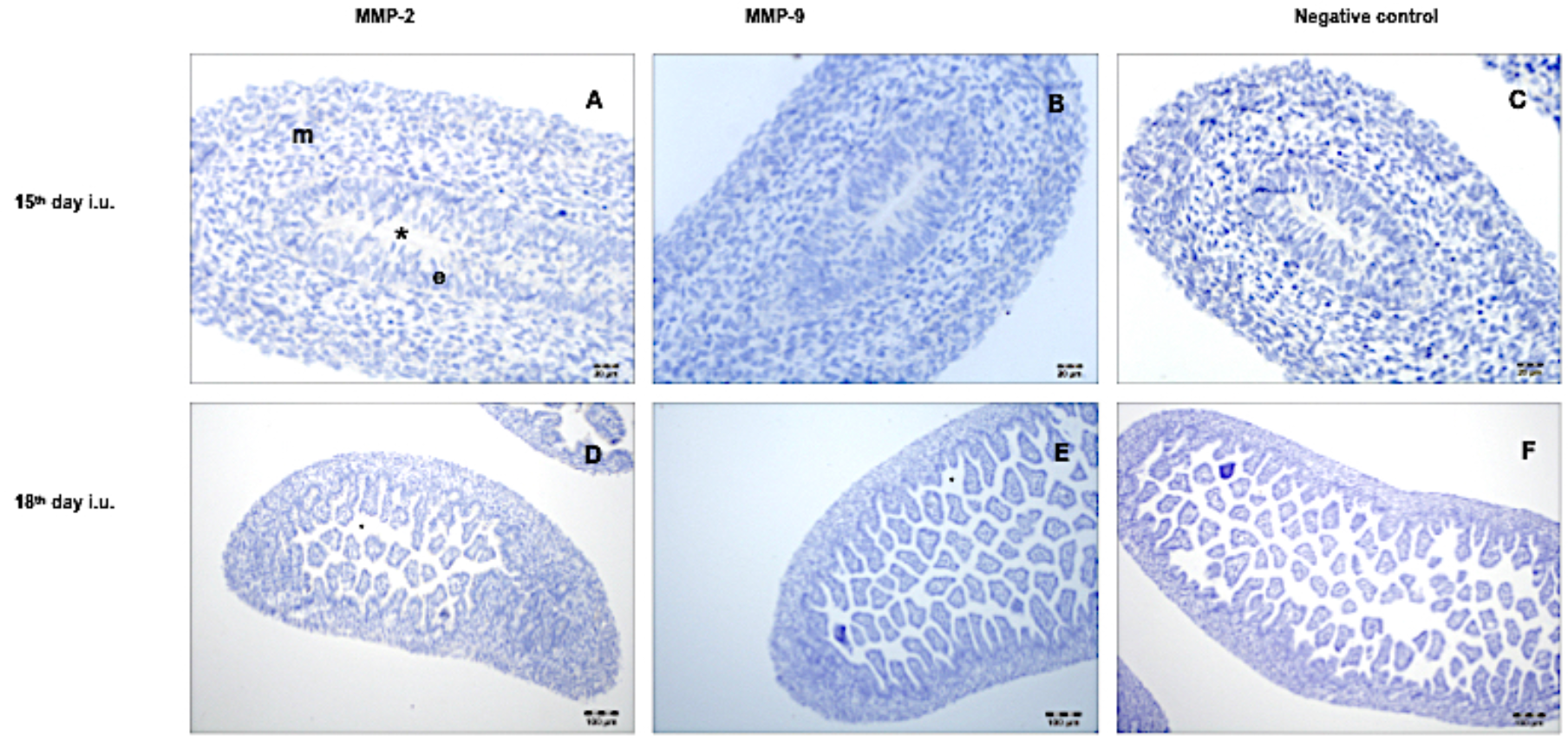
Immunohistochemical staining for MMP-2 and MMP-9 in the small intestine in the times evaluated in the intrauterine life period: 15^th^ and 18^th^ days. Mesenchyme (m); lumem closed (*) and epithelium (e).

### During the postnatal development of the small intestine only MMP-9 is expressed

**Concerning to zym**ogram results obtained for the 3th and 10th days, (suckling period), 17th (pre-weaning period) and 25 and 32th days (weaning period) demonstrated in the Fig. 1A, it was possible to detect a 90 kDA protein with a proteolytic activity, which corresponds to the molecular weight to MMP-9. Another protein with a proteolytic activity was detected at 72 kDA however, only for samples obtained on the 32nd day.

However, after the use of the phenantroline plus EDTA (Fig 1B) as well as the PMSF (Fig. 1C) which are MMP and Serinase inhibitors, respectively, the results show an inhibitory effect on the proteolytic activity at 90 kDA and 72 kDA levels in all times evaluated. However, the iodocetamina, which is a Cisteinase inhibitor (Figure 1 D), was not able to inhibit proteolytic activity in small intestine samples in any time evaluated.

In the immunohistochemistry the MMP-2 was not detected on the 3rd, 10th, 17th, 25th and 32th days as shown in the Figure 3 from A to E. However, MMP-9 was immunolocalized in the goblet cells cytoplasm, in the villus (arrow - Fig 3 F′), on 3rd and 10th days but not in the next times evaluated. MMP-9 was also detected in the cytoplasm of cells localized in the lamina propria in all times evaluated in the postnatal life as indicated by arrows in the Figure 3 from G′ to J′ images.

**Fig. 3.**
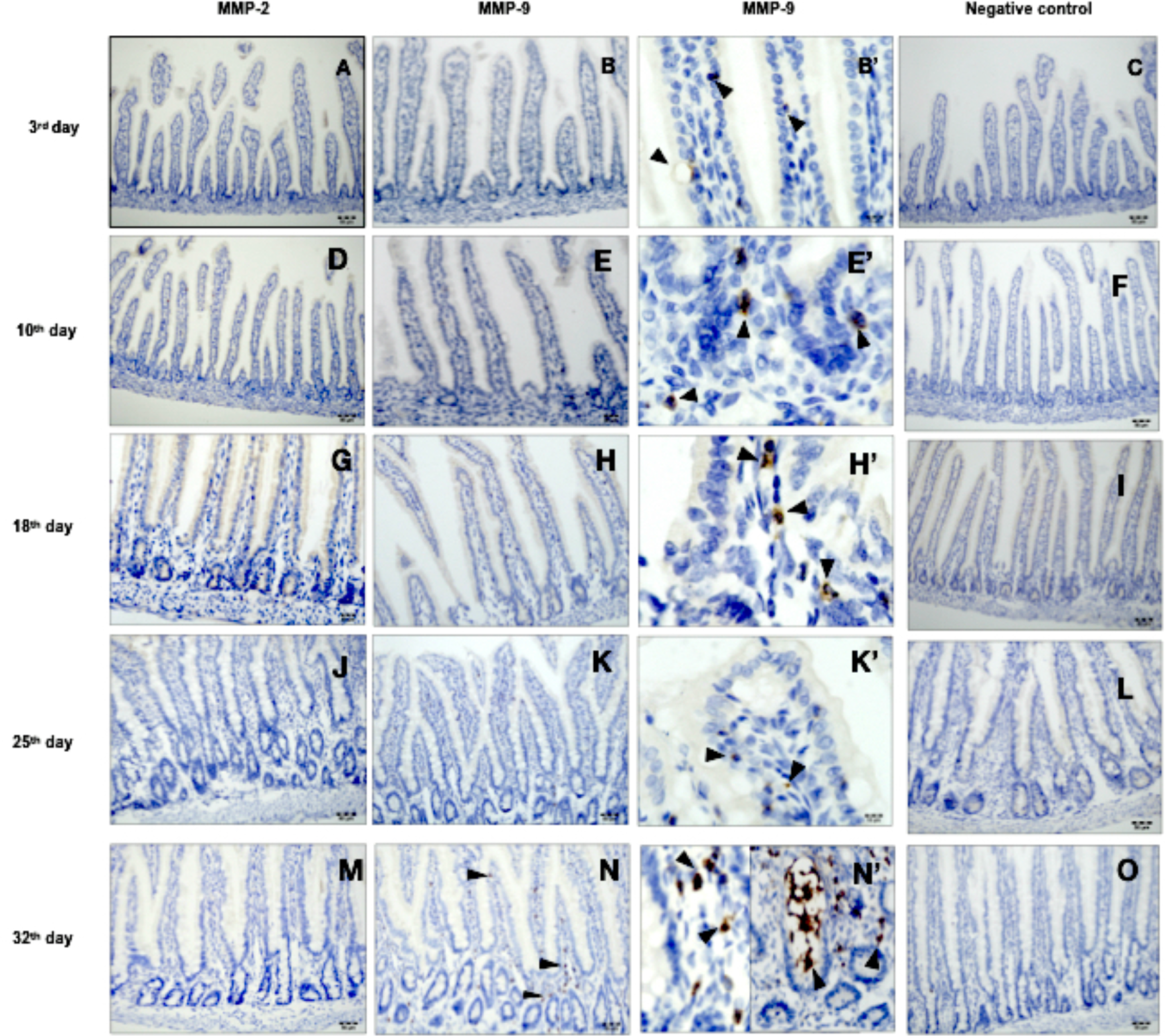
Immunohistochemical staining for MMP-2 and MMP-9 in the mucosa of the small intestine in the times evaluated after birth: 3^rd^, 10^th^ 18^th^ 25^th^ and 32^th^ days. Arrows indicates the positive staining for MMP-9 only in cells present in the lamina propria in the mucosal layer. MMP-2 was not detected after birth.

### The MMP-9 expressed in the lamina propria is produced by the Mast Cell

After have been shown that the most of the positive cells for MMP-9 were in the lamina propria in the 17th, 25th and 32th days after birth, with Masson’s staining was possible to detect that they were mast cells as demonstrated in the Figure 4.

**Fig. 4.**
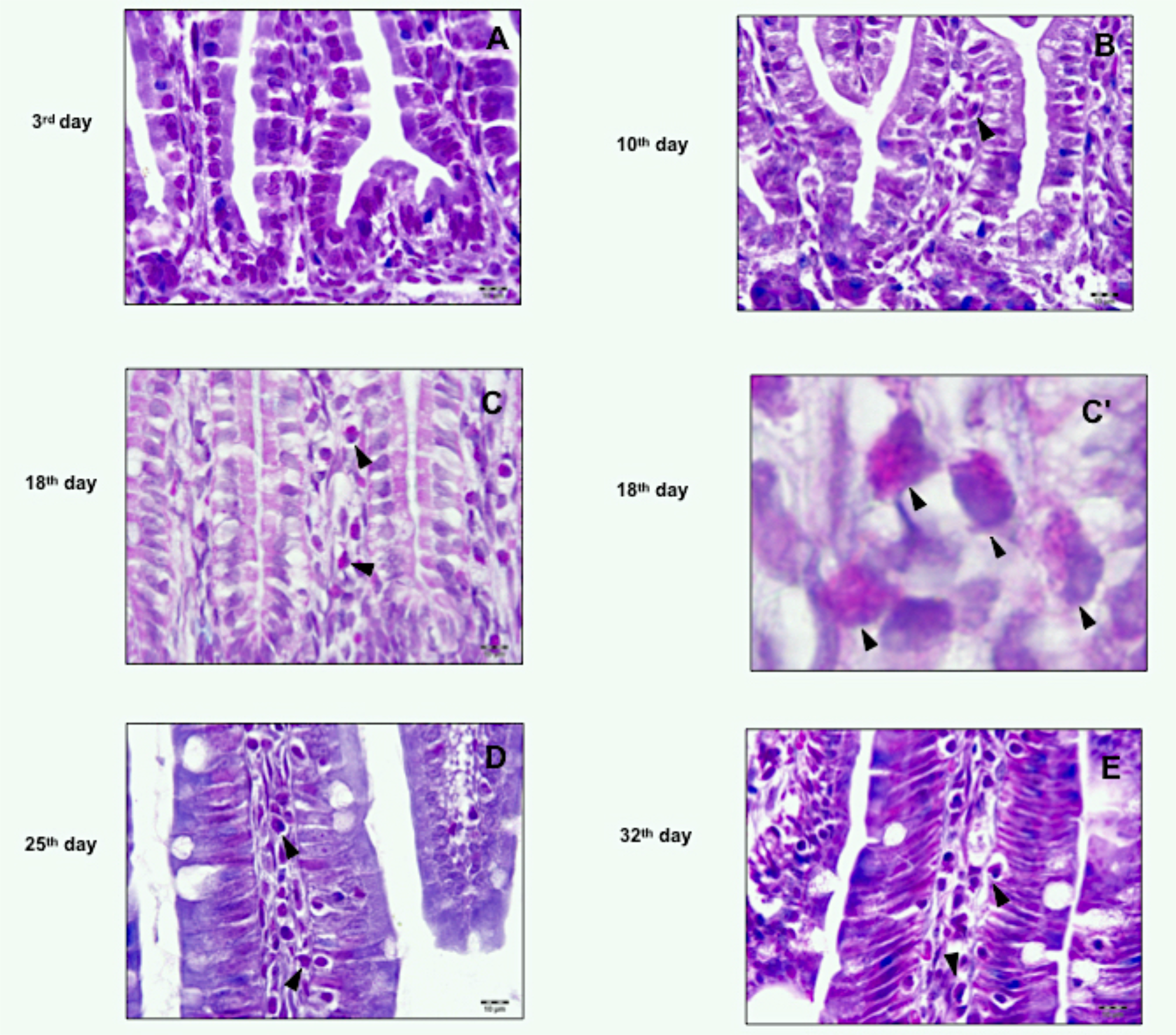
Representative images where the mast cells are indicated by head of arrows in the lamina propria of the small intestine, from the 10_th_ to 32_th_ day, after birth. (C’ is an ampliated image from C). Masson’s thrichome staining.

## DISCUSSION

The small intestine is a vital organ for the homeostasis of body systems because all essential nutrients for the cell metabolism are intaken at small intestine level. In addition, small intestine also acts as a barrier between the body internal and external environment (De Mey and Freund, 2013). However, during all phases of development, the small intestine is under influence of the external factors, mainly light and the food (Soares et al., 2009). In mammalian, the first food intake after birth is milk. This phase is defined as suckling which in rats ends around the 17nd day a.b. After this time, the animal goes to the weaning phase when the food habit becomes diversified, indicating that during the small intestine development there are several adaptations concerning its morphophysiological functions, especially in enzymes expression.

It is known that MMP-2 and MMP-9 are involved in the remodeling and signalization process of the extracellular matrix such as the modulation of the biological growth factors and cell proliferation in normal tissue developmental repair or during the progression of several degenerative diseases (Stamenkovic 2003, Phillips et al., 2017, Nawaz et al., 2018). To perform these functions MMPs are actively expressed by different types of cells like keratinocytes, endothelial cells, fibroblasts, chondroblasts, immune system and Schwann cells (Chattopadhyay and Shubayev 2009, Zambuzzi et al. 2009, Vaillant et al. 2003).

During the development of the organs, the expression of MMPs has been demonstrated in different animal models (Quick et al., 2012; Feng et al;. 2012; Jafari et al., 2010). However, concerning to expression of MMP-2 and MMP-9 during small intestine development of mammalian (from intrauterine to adulthood), the descriptions are scarce. Camargo et al., (2016) demonstrated that MT1-MMP (an MMP anchored in the cell membrane) is expressed by the epithelial cells of small intestine during all phases of the intrauterine and postnatal life. Therefore, we hypothesized that the MMP-2 and MMP-9 also could to be expressed during the morphogenesis of the small intestine considering the intrauterine, suckling and weaning phases which are described as important windows for the intestinal morphogenesis. In special, the presence of the MT1-MMP suggest a possible expression of the MMP-2 because its activation only occurs in the presence of the MT1-MMP (Murphy et al., 1999).

Contrary to MT1-MMP reported before (Camargo et al, 2016), our results demonstrated no expression of the MMP-2 and MMP-9 by epithelial cells of the small intestine during the development in the intrauterine life. This finding indicates that these gelatinases (MMP-2 and MMP-9) may not have any relevant role in this fetal phase of the rat small intestine development. However, after birth although a gelatin band degradation corresponding to the MMP-2 was present in the zymogram on the 32nd day, it was not detected by immunohistochemistry in any of phases evaluated (suckling, pre-weaning, and weaning). This results suggest that on the 32nd day that gelatin band degradation may be not MMP-2. Taken together, the results from the postnatal and the intrauterine life indicates that MMP-2 is not present in any phase of development of small intestine, and in addition to that described by Camargo et.al., (2016), it alowed us to suggest that the MT1-MMP may have the main role during the remodeling of the epithelial layer of the small intestine.

The MMP-9 expression was detected in the zymogram at all times evaluated and also by immunohistochemistry method in the goblet cells (in some times evaluated) and in the cells localized in the lamina propria but not in the epithelial cells, as we had hypothesized․. However, the use of the MMP and Serinases inhibitors reduced the proteolytic activity at 90 kDA. Additional results showed that, in the lamina propria, mast cells were expressing MMP-9. This cell was distributed in the center of the villus, between the epithelial cells and around of mucosal glands from the 17th day after birth. The presence of MMP-9 in the mast cells and the zymograms and inhibitory test are consistent because the literature shows that the mast cells produces several pharmacological molecules including the Chymase and Tryptase (two serinases), which are major proteins present in the granules of the mast cells (Katunuma & Kido, 1988; Lees et. al, 1994). Chymase and tryptase also act in the degradation of the extracellular matrix components. Chymase of mast cells is also involved in the permeability of the epithelial cells changing the distribution of the tight junction proteins (Scudamore et. al., 1988). Therefore, the results for the inhibitory test using the PHE and PMSF showed a reduction in the band degradation for both inhibitors which suggest that the band present in the PHE test is the serinase and the band present in the PMSF test is MMP-9. This is a plausible and interesting interpretation obtained in this investigation, once that the mast cells may express both MMPs and Serinases.

In addition, the presence of mast cells in the gastrointestinal wall may be implicated in several biological functions, such as allergic reactions to food, smooth muscle contraction and peristalsis, induction of acid secretion, electrolytes and mucines by the epithelial cells, and host defense against microbes like bacteria, viruses, or parasites (Bischoff, 2009). The role of mast cells and its pharmacological molecules on the inflammatory modulation as well as in the adaptative and especially in the innate immunity has been also discussed (Pejler et al., 2010; Douaiher et al 2014; Cardamone et al., 2016; Pastwińska et al 2017; Xu et al., 2018). The activation of the pro-MMP-9 was demonstrated in vitro in the presence of the mouse mast cell protease-4 (mMCP-4) obtained from bone marrow-derived cultured (Tchougounova et. al., 2005; Lin et al., 2011).

In spite of the knowledge existing about the mast cell and its presence in several organs including small intestine, until the current moment we could not find any report about the relationship between the mast cells and the phases of the small intestine development (suckling, pre-weaning and weaning). Because that mast cells expressing MMP-9 were detected from the 10th day after birth, we suggest that it may be involved, for example, in the remodeling of extracellular matrix in case of the inflammatory process, as observed in the others immunological cells such as macrophage (Butoi et., al 2016) however, this still needs to be confirmed during the small intestine development in further investigations.

In conclusion, our results showed that MMP-2 and MMP-9 were not expressed in the epithelial cells during the intrauterine and post natal development of the small intestine however, MMP-9 was expressed in the mast cell present in the lamina propria after birth indicating its involvement in the development of the small intestine innate immunity.

## Acknowledgements

The authors thank to Dr Danilo Massuia Rocha for performing English review, to Histology laboratory and NAEVI of UEPG for metodological and animal support respectivelly.

## Author contributions

Conceptualization, Methodology, Funding acquisition and Writing: Gomes, JR. Development, Execution of Procedures, Results and Analisys: Reis, CA.

